# Crystal structure determination of the antipsychotic drug of olanzapine form III

**DOI:** 10.1101/2024.05.15.594141

**Authors:** Goulielmina Anyfanti, Elena Husanu, Iryna Andrusenko, Danilo Marchetti, Mauro Gemmi

## Abstract

Olanzapine, an antipsychotic drug, is well known for its complex polymorphism. Although widely investigated, the crystal structure of one of its anhydrous polymorphs, form III, is still unknown. Its appearance, always in concomitance with form II and I, and the impossibility of isolating it from that mixture, has prevented its structure determination so far. The scenario has changed with the emerging field of 3D electron diffraction (3D ED) technique and its great advantages in the characterization of polyphasic mixture of nanosized crystals. In this study we show how the application of 3D ED allows the ab-initio structure determination and dynamical refinement of this elusive crystal structure unknown for more than 20 years. Olanzapine form III is monoclinic and shows a similar but shifted packing with respect to form II. It is remarkably different from the lowest energy structures predicted by the energy minimization algorithms of crystal structure prediction.

---

Olanzapine (OLZP) (fig. 1), a thienobenzodiazepine derivative, is a second-generation antipsychotic mostly used for the treatment of schizophrenia and bipolar disorder due to its antagonist action for multiple neurotransmitter receptor sites.^[1]^

**Figure 1.**
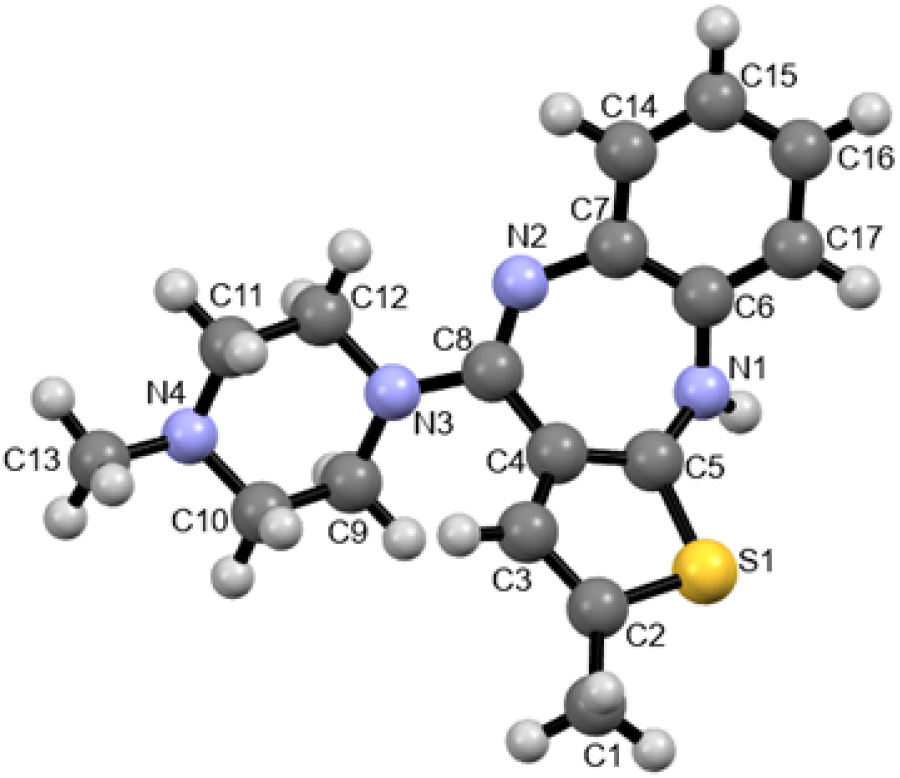
Olanzapine molecular structure with the relative label scheme.

Its ability to assume different solid-state phases forming anhydrous, hydrated, and solvated polymorphs^[2–10]^ represents the reason of the intense research work developed around its polymorphism. When it comes to the pharmaceutical industry, the control of the possible solid forms and, therefore, of the physicochemical properties such as bioavailability, solubility and stability is the basic requirement in drug development^[11–15]^ and a key point of patenting.^[16]^

OLZP counts more than 60 polymorphs, with only four being known as anhydrous phases. Three of these, form I, II and III, have been discovered during the development of the marketed pharmaceutical Zyprexa at Lilly research laboratories^[7]^, and resulted in a patent which dates 1993.^[17]^ Form I has been confirmed as the thermodynamically stable and selected for the commercial development into Zyprexa.^[7]^ It is the only form that was crystallized as a single phase by direct crystallization from solvents that do not form solvate with OLZP. Forms II and III can be crystallized as desolvation products of certain OLZP solvates^[9]^ but, until now, could not be isolated as pure polycrystalline phases. Both forms I and II can be grown in single crystals large enough for single crystal X-ray diffraction, so their structure has been determined.^[2,4,5]^ All the efforts of different research groups to obtain OLZP form III in pure form failed to the best of our knowledge and the concomitant appearance of phase III with phase II and its microcrystalline character, has prevented its structure determination so far. A possible structure has been proposed via crystal structure prediction (CSP) techniques,^[5,18]^ but a trial to fit powder X-ray diffraction with a two phases Pawley-type refinement was not full satisfactory, since some peaks belonging to phase III were not properly modeled. Both phase I and II, as well as the proposed structural model of phase III, which is an alternative layer stacking of phase II, are based on a “dispersion” dimer of opposite enantiomer pairs related by a center of symmetry identified as the supramolecular construct SC_0_ by Bhardwaj et al.^[5]^ The dimer is held by dispersion forces and the two enantiomers have exposed on the outer part hydrogen bonding donors and acceptors responsible for the crystal structure packing. Interestingly the CSP study that delivered the only available structural model of phase III proposed also, among others, some OLPZ structures in which the SC_0_ building was absent. This triggered a dedicated search of new OLPZ polymorphs through non – conventional crystallization methods like, polymer based molecular dispersion, which resulted in the discovery of the new anhydrous form IV.^[6]^ In this scenario the crystal structure of phase III is, for the moment, the only missing tile in the puzzle of anhydrous OLZP polymorphism.

Although crystalline structural changes are generally monitored through single crystal and powder X-ray diffraction (PXRD) analysis, gaining an overview of the molecular arrangement and their relationship with bulk properties, the limitations given by the nature of OLZP phase III challenge the two complementary techniques for its structure determination. In recent years many studies have shown the effectiveness of single crystal electron diffraction, known as 3D ED or MicroED,^[19]^ in elucidating the structure solution of organic nanocrystals of pharmaceutical interest.^[20–25]^ For these reasons, and for the importance of the structure dilemma of OLPZ phase III we decided to examine the powder mixture of OLZP forms I, II and III with 3D ED and we surprisingly have been able to solve the crystal structure of the latter ab-initio. Dynamical refinements led us on the one hand, to the final model of form III that showed a different packing of layers with respect to form II. On the other hand, the structure of OLPZ form III was solved in P2_1_/c and not Pbca, differently from the predicted model (named A162 by Bhardwaj et al.^[5]^).

Starting from a pure OLZP form I sample (see the Le Bail^[26]^ fit in fig. S1) a mixture of OLZP forms I, II and III with a higher content of the latter was obtained using a slightly modified method of crystallization with respect to the one published by Reutzel-Edens group (see supporting information).^[2]^ The characteristic powder pattern and differential scanning calorimetry (DSC) thermograms (fig. S2 and S3) of the mixture coincide with what observed by Bhardwaj et al. ^[5]^ The sample, under the transmission electron microscope (TEM), revealed the presence of micro to nanosized crystals. Although crystal habits prediction studies ^[27,28]^ have described how olanzapine crystals of the same polymorph can adopt different morphologies depending on the crystallization solvent used^[28]^, all the grains observed under the TEM show an irregular shape and any distinction based on the habit of the nanosized crystal was impossible (fig. S4).

3D ED data were collected over 32 crystals and their indexing confirmed the presence of three different polymorphs. A first group of crystals which could be indexed as form II, a second very rare group which could be indexed as form I, and a third one which could be indexed with a monoclinic cell never reported before. We considered the latter as representative of form III. While for form I we determined only the unit cell parameters, the structure of both forms II and III could be solved ab-initio. Table 1 shows the unit cell parameters of the three forms.

**Table 1.** Unit cell parameters of forms III, II and I from the data collected with 3DED at room temperature.

| Polymorph | III | II | I |
| --- | --- | --- | --- |
| a (Å) | 10.708(3) | 9.83(5) | 10.33(5) |
| b (Å) | 16.476(4) | 16.60(7) | 15.262(3) |
| c (Å) | 10.065(4) | 9.943(4) | 10.52(1) |
| α (°) | 90 | 90 | 90 |
| β (°) | 110.43(2) | 98.08(4) | 100.5(6) |
| γ (°) | 90 | 90 | 90 |
| V (Å<sup>3</sup>) | 1664 | 1607 | 1630 |

The crystallographic data along with the results of the dynamical and kinematical refinements for phases III and II are listed in table S1. The correspondent images of the hk0, h0l and 0kl reciprocal space sections are shown in figure S5. The structure of form II was solved and kinematically refined and it corresponds to the one reported in the literature, while form III was new and different from the predicted model A162. It crystallizes in the monoclinic space group, P2_1_/c as form II, but with a larger monoclinic angle 110° instead of 98°.

The quality of the collected 3D ED data prompted us to refine form III structural model considering the dynamical diffraction theory.^[29,30]^ Dynamical refinement takes into account the multiple scattering effects delivering more precise structures and fitting much better the experimental intensities. Still not a diffuse practice in the case of unknown organic structures, the use of dynamical refinement results in a drop of the residual R-value which usually reduces to a half with respect to kinematical refinement. In our case the solved structure of form III was dynamically refined against two data sets at a resolution of 1 Å reaching a final R-value of 12.23%. The thickness variation during crystal rotation was modeled with the thick model wedge approximation.^[31]^ Atomic displacement parameters (ADPs) of all non-hydrogen atoms were refined anisotropically. Atoms in a similar chemical environment were restraint to have similar anisotropic ADPs. For instance, the anisotropic ADPs of C15, C16 and C17 carbon atoms were kept similar to that of C14. The 3D ED data quality is sensitive to the presence of hydrogens.^[32]^ While some of them can be already identified in the difference Fourier map before the refinement (fig.S6), neglecting the hydrogens significantly worsen the refinement.

This can be checked, not only by the R-value, that drops from 15.77% (R_obs_) to 12.23% (R_obs_), but also by the difference Fourier map that, once aligned and scaled to the same isosurface value using Vesta^[33]^, is much less noisy after the hydrogen addition than before (fig. S7).

A Le Bail fit of the PXRD pattern (fig. S8) collected on the sample confirmed the mixture of the three forms, while a Rietveld refinement (fig.2, R_obs_ = 3.11%, wR_obs_ = 4.26%) further assures that the structural model of form III derived from 3D ED is correct (fig. 2). The Rietveld also states that form III is the major phase with a volume fraction of 68% followed by form II, 25.8%, and an almost negligible amount of form I, 6.2%, in agreement with previous observations.^[5]^

**Figure 2.**
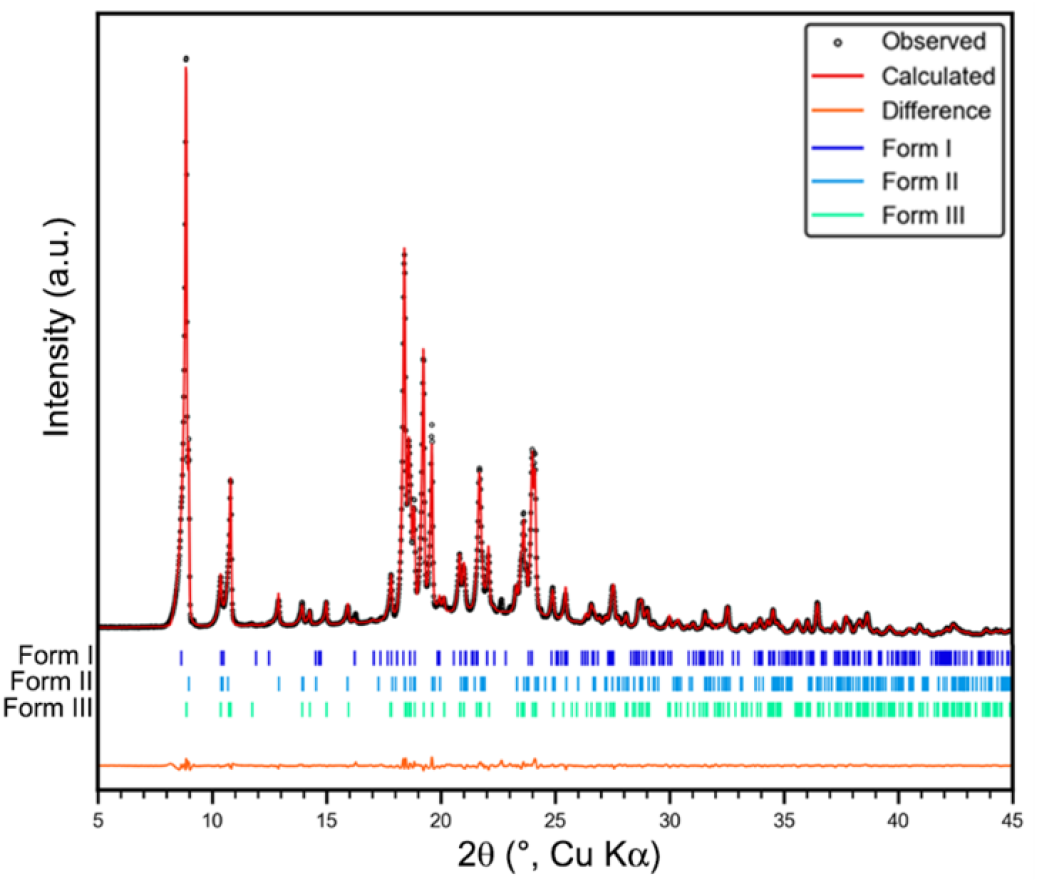
Three-phases Rietveld plot of the multiphase powder sample

The characteristic building block of phase III is the same face-to-face centrosymmetric dimer motif known as the supramolecular construct SC_0_ ^[5]^ (fig. 3) encountered in forms I and II and also in the predicted structure of form III, the orthorhombic A162. It consists of head-to-tail enantiomers bound with each other by purely dispersion interactions. Each enantiomer assumes the same conformation observed and described by Reutzel-Edens ^[2]^ where the piperazine ring in its chair conformation is almost coplanar with the puckered diazepine ring. Each SC_0_ dimer interacts with the adjacent ones through dispersive interactions and a unique hydrogen bond network (fig. 4a). Inside this network we can distinguish a NH···N interaction between N1, as donor, and the neighboring N2 (x, 3/2-y, −1/2+z), as hydrogen bond acceptor. Another weaker intermolecular interaction that contributes to the packing stability of OLZP form III is a bifurcated contact between C17-H, as donor, and N2 (x, 3/2-y, −1/2+z) or C14 (x, 3/2-y, −1/2+z) as acceptor (CH···π). (fig.4b). As in form II also in phase III the bonding network drives the formation of corrugated planes parallel to (100) that if looked along the c direction exhibit a wavy shape (fig. 4c). Up to now we have dealt with the arrangement of the intermolecular interactions of form III by considering the number of atom-atom contacts that have the greatest effect on the stability of the crystal, for instance the hydrogen bonds. The whole-to-molecule approach adopted in the calculation of the Hirshfeld isosurfaces^[34–36]^ gives, instead, a global visualization of the type of short contacts involved in the crystal packing. Figure 5 compares the Hirshfeld surfaces calculated for forms II and III. The red areas highlight the close contacts of the molecule with the surroundings and in both phases, they coincide with what has already been described. The sulfur atom of the thiophene ring also forms close contact with neighboring molecules. The relative contribution of each intermolecular interaction on each surface is shown in the fingerprint graphs^[37,38]^ of the two surfaces in figure S9 as well as in table S2 In all forms the dispersion dimer is the unit that, repeated along a plane direction (bc in form II and III and ac in A162), forms 2D layers. The main difference between forms II, III and the predicted form A162 arises in the way the 2D layers are packed in the third dimension. To better understand and display this difference we first compared the structure of form II with our model of form III.

**Figure 3.**
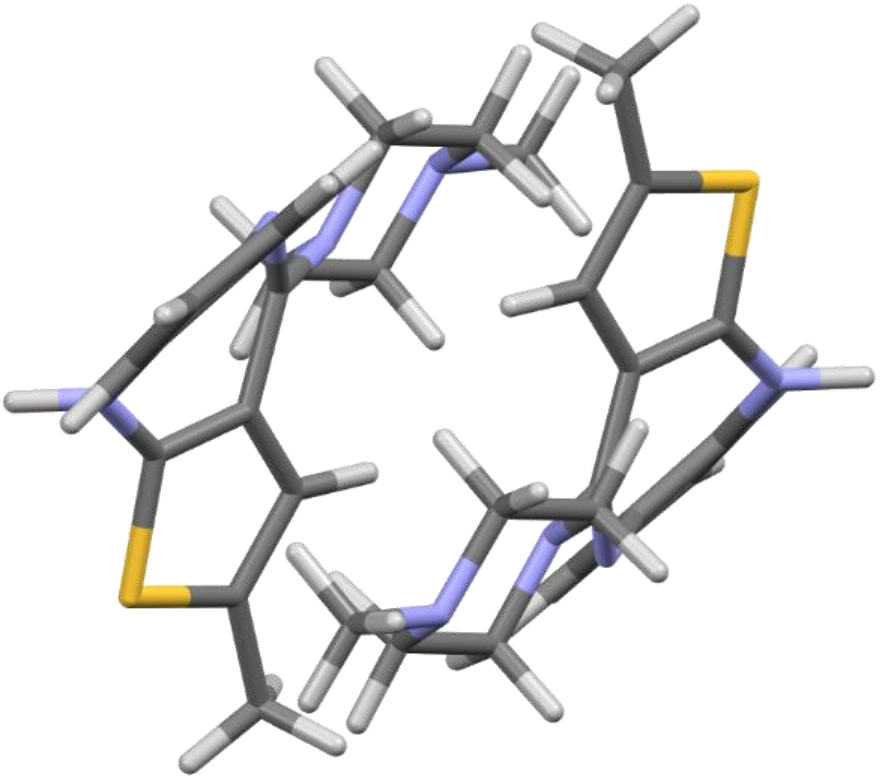
Head-to-tail enantiomers form the centrosymmetric building block SC_0_ of OLZP form III.

**Figure 4.**
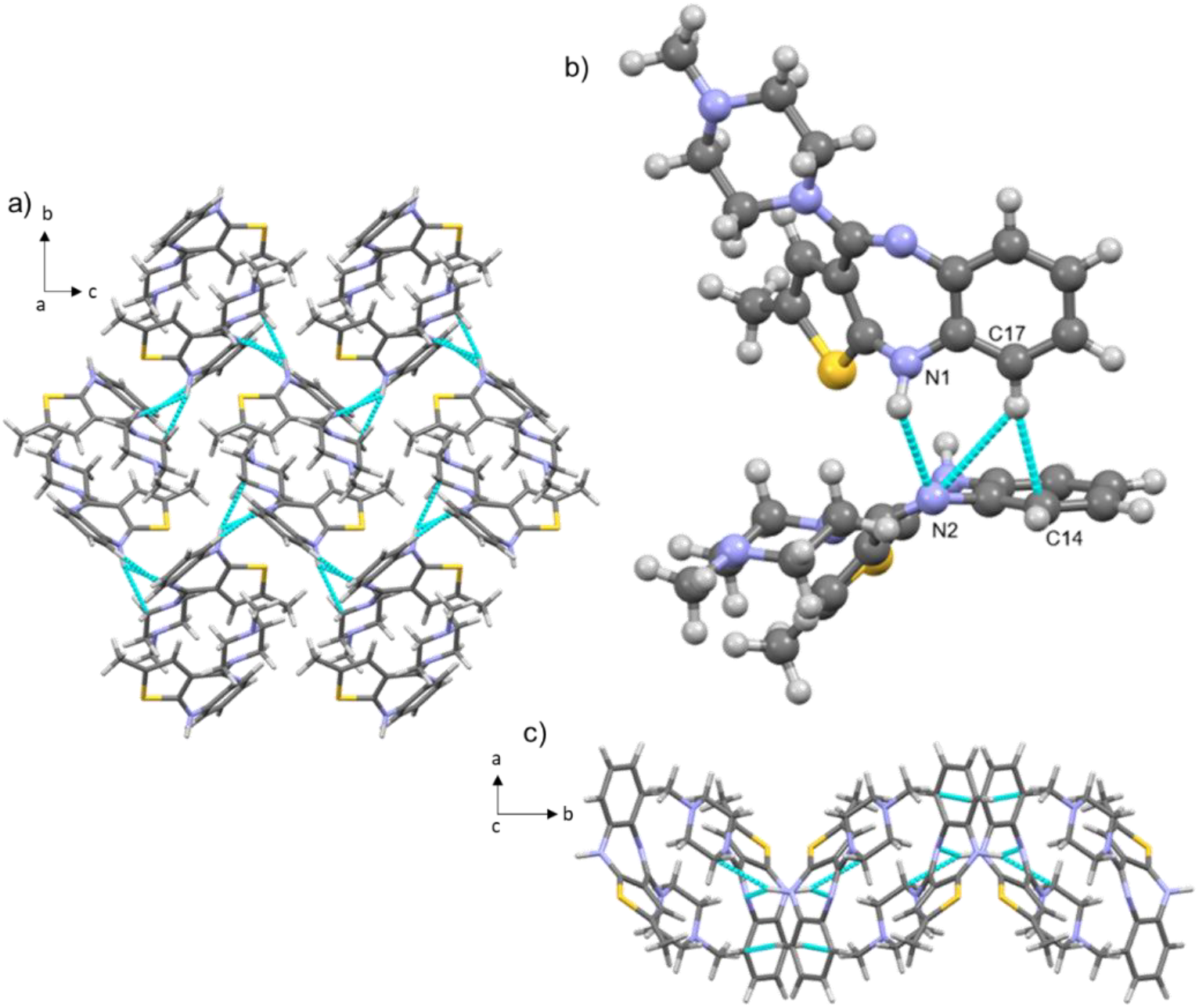
One 2D layer of OLZP form III viewed along a-axis with the relative intermolecular interactions among adjacent SC_0_ dimers (a), closer view of the intermolecular contacts between two adjacent molecules (b) and one 2D corrugated plane of OLZP along the c-axis with the relative intermolecular contact network (c).

**Figure 5.**
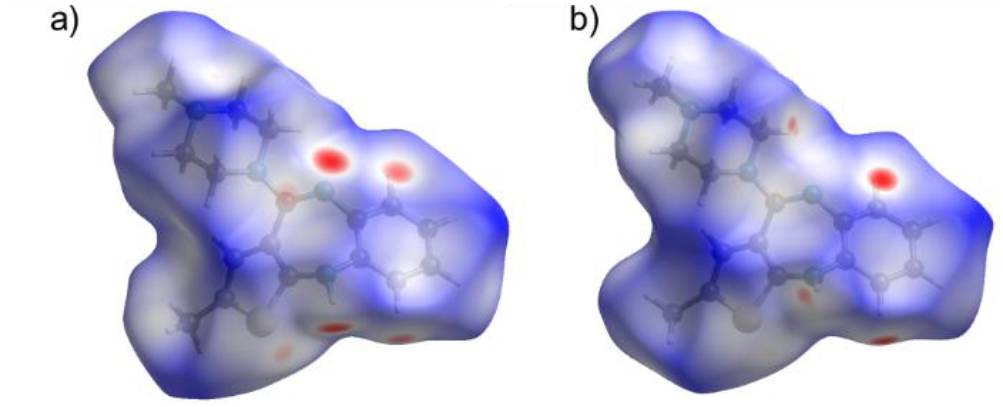
Hirshfeld surface of an OLZP molecule in forms II (a) and III (b)

Figure 6a shows the stacking of three consecutive layers viewed along the c-axis for forms II and III, colored alternatively in green and pink. Looking at the structure perpendicularly to these layers and taking as a reference the orientation of the dimers viewed along the a-axis of form II, the difference in the stacking can be clearly highlighted. In form II all the dimers of the different layers perfectly superimpose (only the pink ones are visible) (fig. 6b), while in form III we can observe a clear shift of the green and pink levels along c (fig. 6c) which is responsible of the increase in the β angle of nearly 12° between the two structures.

**Figure 6.**
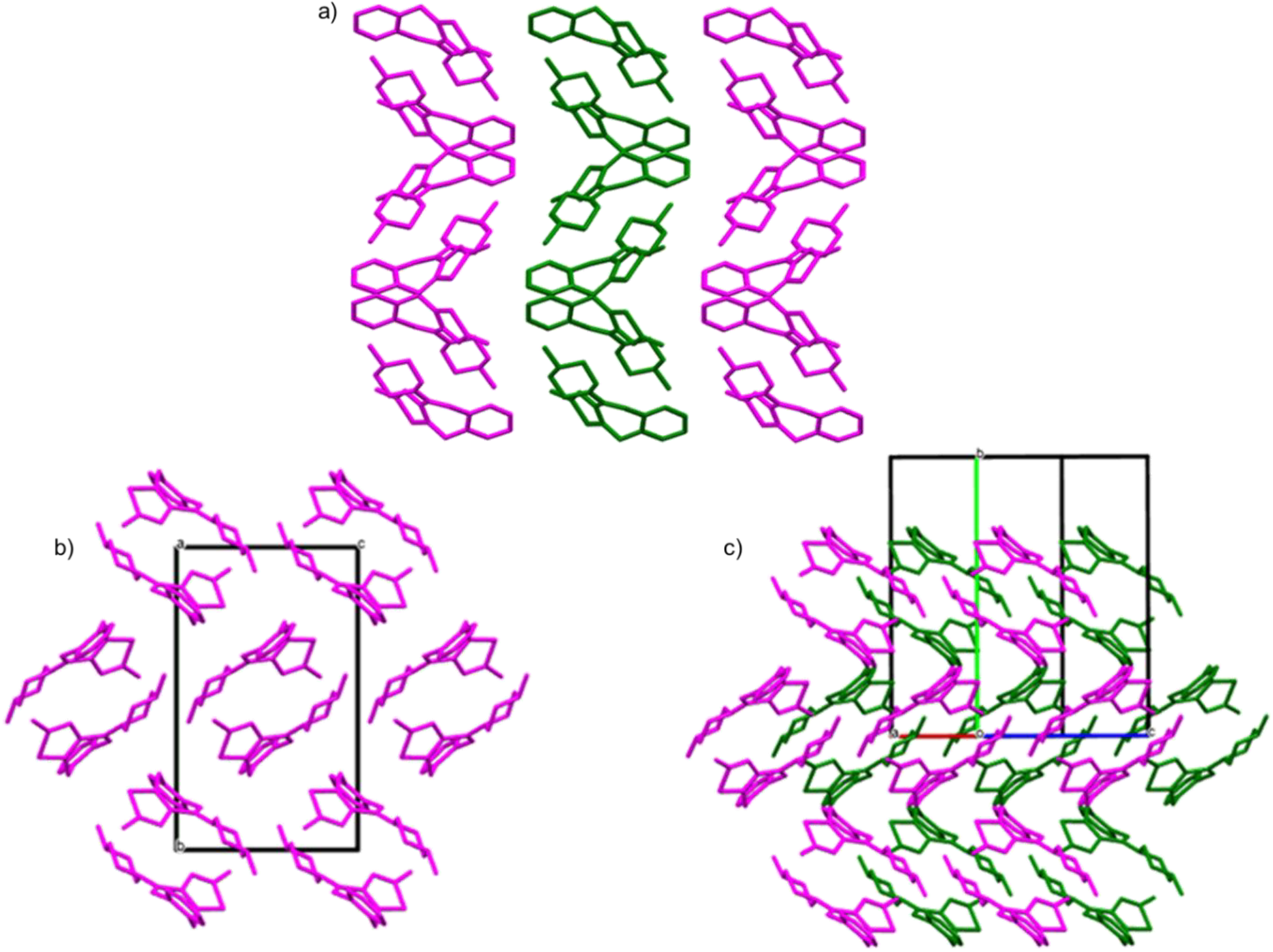
Packing of three layers in forms II and III respectively viewed along the relative c-axis, the central layer is in green for the visualization of the different packing (a); the two structures are rotated and viewed along the a-axis in form II (b) and along the equivalent crystallographic axis in the found monoclinic structure of phase III where the shift of the pink layers is evident (c).

The displacement along the third dimension is the main difference claimed also by Bhardwaj in A162, but, if in form III there is just a shift along one direction every second layer, in A162 the shift is obtained through a c glide perpendicular to b, which causes the exchange of the enantiomers every second layer. For a better understanding see figure S10.

A careful screening of the CSP structures of OLZP obtained in Bhardwaj et al. study and kindly provided by the authors lead to take into consideration one predicted structure, identified in the original study as UNOGIN_eq_125 as the most similar to our form III model. However, this predicted structure does not belong to the lowest energy structures predicted on the crystal energy landscape. The root-mean-square deviation (RMSD) obtained from the overlay of 90 molecules of the two crystal structures in the Crystal Similarity Tool^[39]^ is 0.329 Å (fig. S11). Characterized by the same SC_0_ dimeric building block, UNOGIN_eq125 packs in the same P2_1_/c space group of form III but the unit cell parameters are slightly distorted with respect to form III (table S3), while the calculated powder pattern differs from the one of phase III at resolution higher than 5 Å (above 18° in 2θ) as it can be seen in figure S12. Apart from these differences, we can say that UNOGIN_eq125 is a good approximation of form III, but unfortunately it seems that the energy of the system is quite sensitive to small distortions, and this has hampered CSP to find the correct minimum.

In conclusion we can claim to have successfully found the missing tile to the OLZP puzzle determining the anhydrous form III by mean of 3D ED. This crystallizes in the monoclinic P2_1_/c space group and the results were confirmed by a successful Rietveld refinement of the OLZP starting phase mixture. The form III of OZP is characterized by the same dispersion dimer seen in phases I and II. Although very similar to form II, it shows a slightly different packing with a characteristic shift every second 2D layer. Remarkably the structure of form III differs from all the predicted crystal structures by CSP except one that results close but not identical, although does not belong to the group of lowest energy. Our results confirm once again that the use of 3D ED for characterizing elusive pharmaceutical phases is a winning solution able to solve crystallographic problems that have been lying unsolved for many years.

## Supporting Information

The authors have cited additional references within the Supporting Information.^[31 - 49]^

### Experimental Section

OLZP was purchased from TCI, dichloromethane was purchased from Sigma. A determined quantity of OLZP as received was completely dissolved in dichloromethane. A yellow liquid solution was immediately obtained, filtered with a 0.45 μm PTFE filter attached to a 5 ml syringe in a petri dish and let to evaporate under the hood. To assure the complete removal of the solvent, the powder obtained was transferred in a BUCHI oven at 40 °C under reduced pressure.

PXRD patterns of the sample packed in 0.5 mm borosilicate glass capillary were recorded in transmission mode at room temperature on a STOE STADI-P diffractometer (STOE, Darmstadt, Germany) equipped with a Ge (111) monochromator and a Mythen2 1K detector from DECTRIS, using Cu-Kα1 radiation (λ=1.5406 Å). The data acquisition and first analysis were processed with WinXPOW software (STOE). DSC measurements were performed with a Mettler Toledo DSC1 analyzer in a N2 flow and a heating rate of 10°C/min.

Le Bail intensities extraction and Rietveld refinements of the powder mixture were performed in Jana2020.^[30]^

Microcrystals from the powder mixture were deposited on 300-meshed copper grids coated with carbon. Electron diffraction experiments were performed on a Zeiss Libra 120 TEM operating at 120 kV and equipped with an LaB_6_ thermionic source and an in-column omega filter. Data were obtained in continuous rotation mode as reported in the paper by Gemmi & Lanza.^[40]^ During the data collection the crystal position was tracked in high magnification scanning transmission electron microscopy (STEM) imaging as proposed by Yang et al.^[41]^.Diffraction patterns were collected in nanodiffraction mode with a parallel beam of 150 nm obtained using a 5 μm condenser aperture. 3D ED diffraction data were recorded using an ASI Timepix detector.^[42]^

Peak indexing and intensity integration were performed using Pets2.^[29]^ Space group determination and structure solution for phase III were obtained with Superflip^[43]^ in Jana2020.^[30]^ Structure of form II was solved by directs methods in Sir2019.^[44]^ Dynamical refinements for form III were performed against two data set of two different crystals at a resolution of 1 Å. The lattice parameters of crystal 2, therefore, were fixed to be the same as those of crystal 1. The data sets collected from crystals 1 and 2 were kept in two data blocks (Block 1 and Block 2 respectively) and a combined refinement against the two blocks was performed in Jana2020.^[30]^ The processed data set for continuous rotation was refined with the thick model wedge^[31]^. The orientation of each patterns was optimized and the ADPs were refined as anisotropic (ADPs of hydrogen were kept anisotropic with the riding model on). In the thick model wedge the crystal is assumed to be wedge shaped. Structure of form II was kinematically refined with SHELXL^[45]^ in Olex2 GUI.^[46]^

Molecular graphics were created with Mercury 4.0.^[47]^ The repository of predicted crystal structure for OLZP was provided by the authors and a screen was performed using the Crystal Packing Similarities (CPS) module provided in Mercury software.

The Hirshfeld surfaces were generated using the CrystalExplorer software (version 21.5) at very high resolution.^[48]^

#### Le Bail intensity extraction of OLZP as purchased

**Figure S1.**
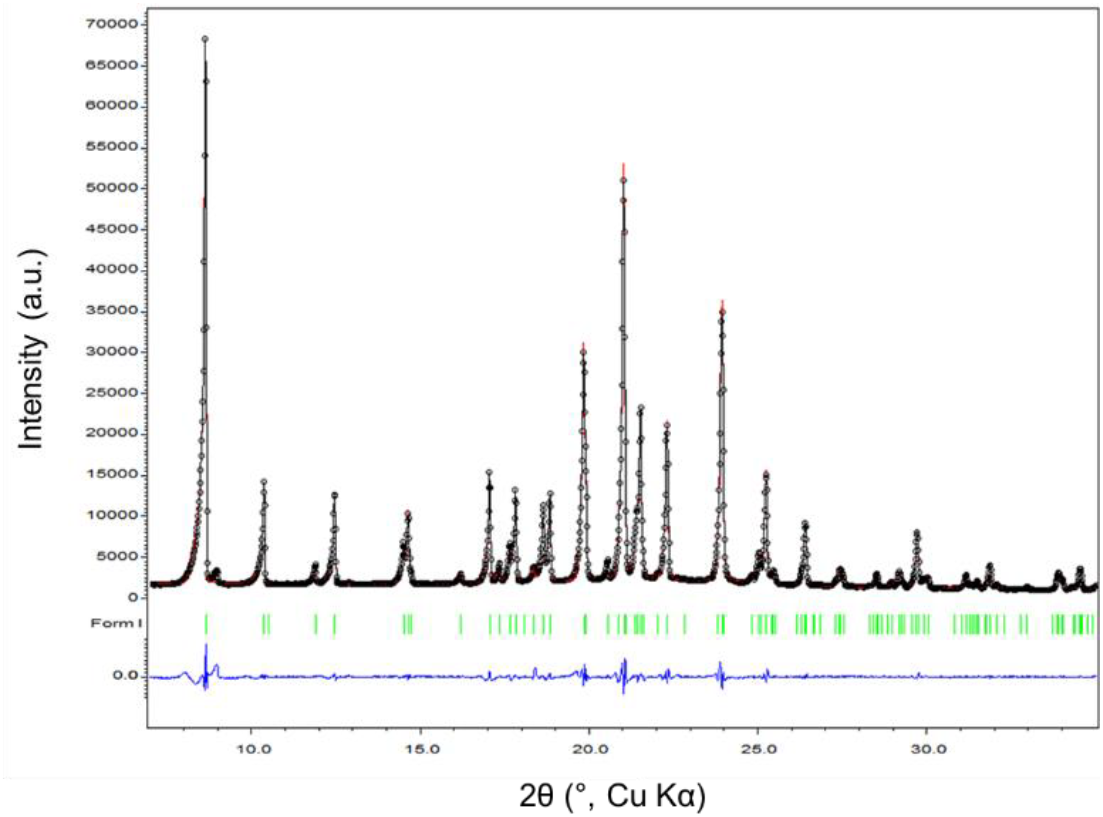
Le Bail fit of OZPN purchased from TCI®. The refined cell dimensions taken from form I ^[3]^ are a = 10. 39821(14) (Å), b = 14. 86624(14) (Å), c = 10. 57299(12) (Å), β = 100. 6391(19)°, R_obs_ = 3.12%, wR_obs_ = 4.72%.

#### PXRD and DSC of the multiphase mixture

**Figure S2.**
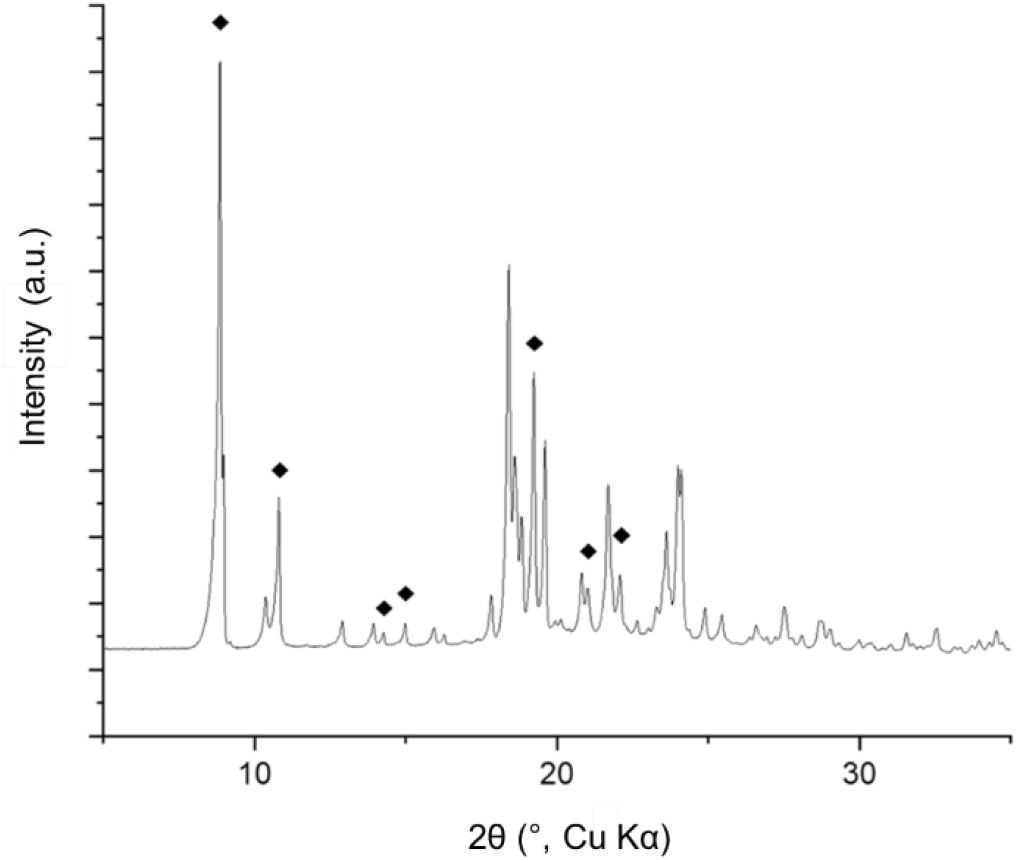
PXRD pattern of the multiphase mixture obtained by recrystallisation of purchased OLZP in dichloromethane. The peaks that testify the presence of multiple phases are signed with the black rhomboidal symbol and correspond to what observed by Bhardwaj et al.^[5]^

**Figure S3.**
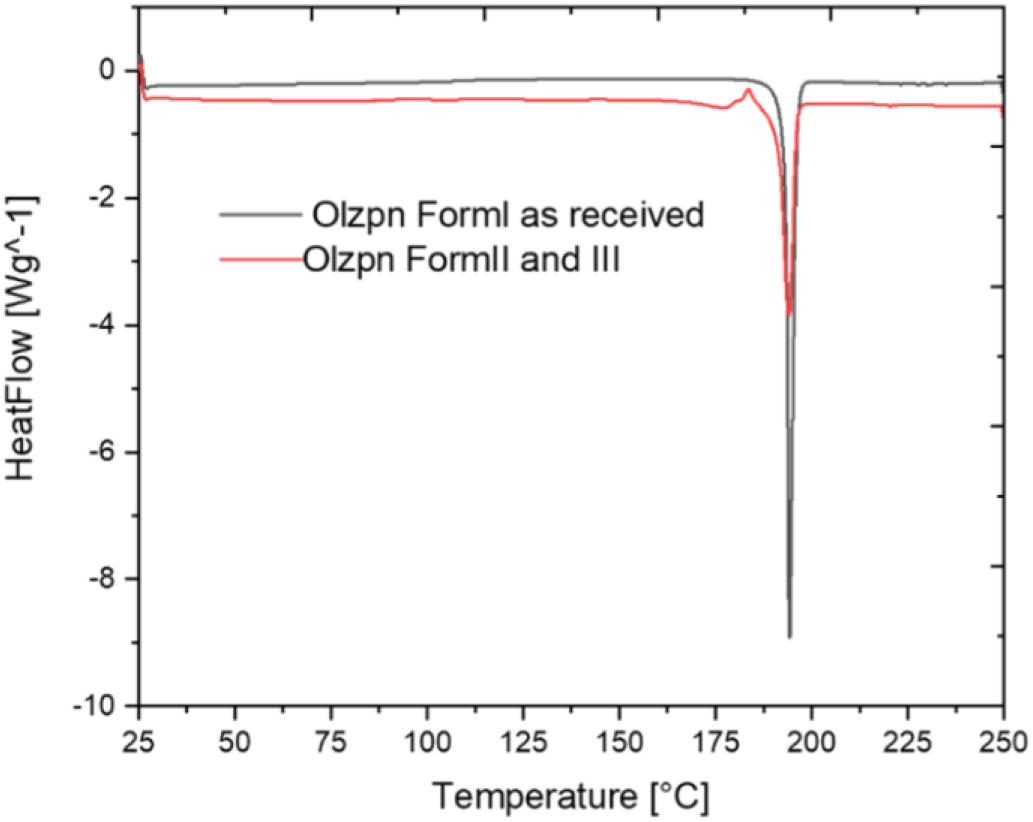
DSC thermograms of powder OLZP form I as received (black line) and the mixture of form II and III (red line) obtained at a flow rate of 10°C/min between 25°C and 250°C upon heating.

#### Crystallographic images and information of forms III, II and I of OLZP

**Figure S4.**
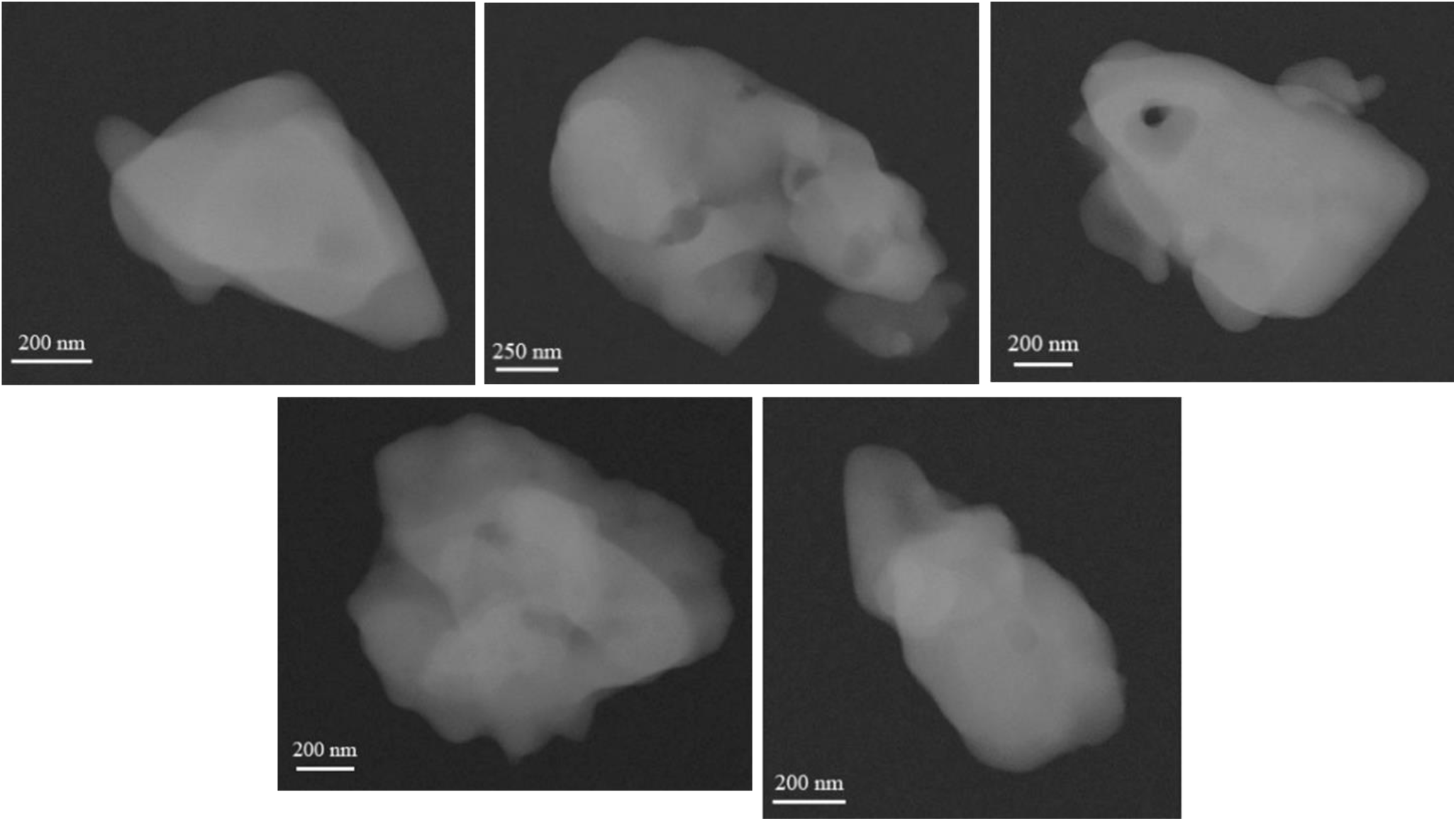
STEM images of OLZP nanocrystals characteristic of mixture form II and III and a low content of form I.

**Table S1.** Crystallographic information for OLZP form III (dynamically refined) and form II (kinematically refined).

| Polymorph | III | II |
| --- | --- | --- |
| Z / Z' | 4/1 | 4/1 |
| Independent reflections | 3084 | 1523 |
| Completeness (%) | 88 | 66.9 |
| I/sig(I) | 4.5 | 4.5 |
| R<sub>int</sub> | 29.60 | 18.28 |
| n° restraints | 269 | 0 |
| GoodF | 2.23 | 1.4 |
| R<sub>1</sub> (%) | 12.23 | 20.61 |
| wR<sub>2</sub> (%) | 12.26 | 51.91 |

**Figure S5.**
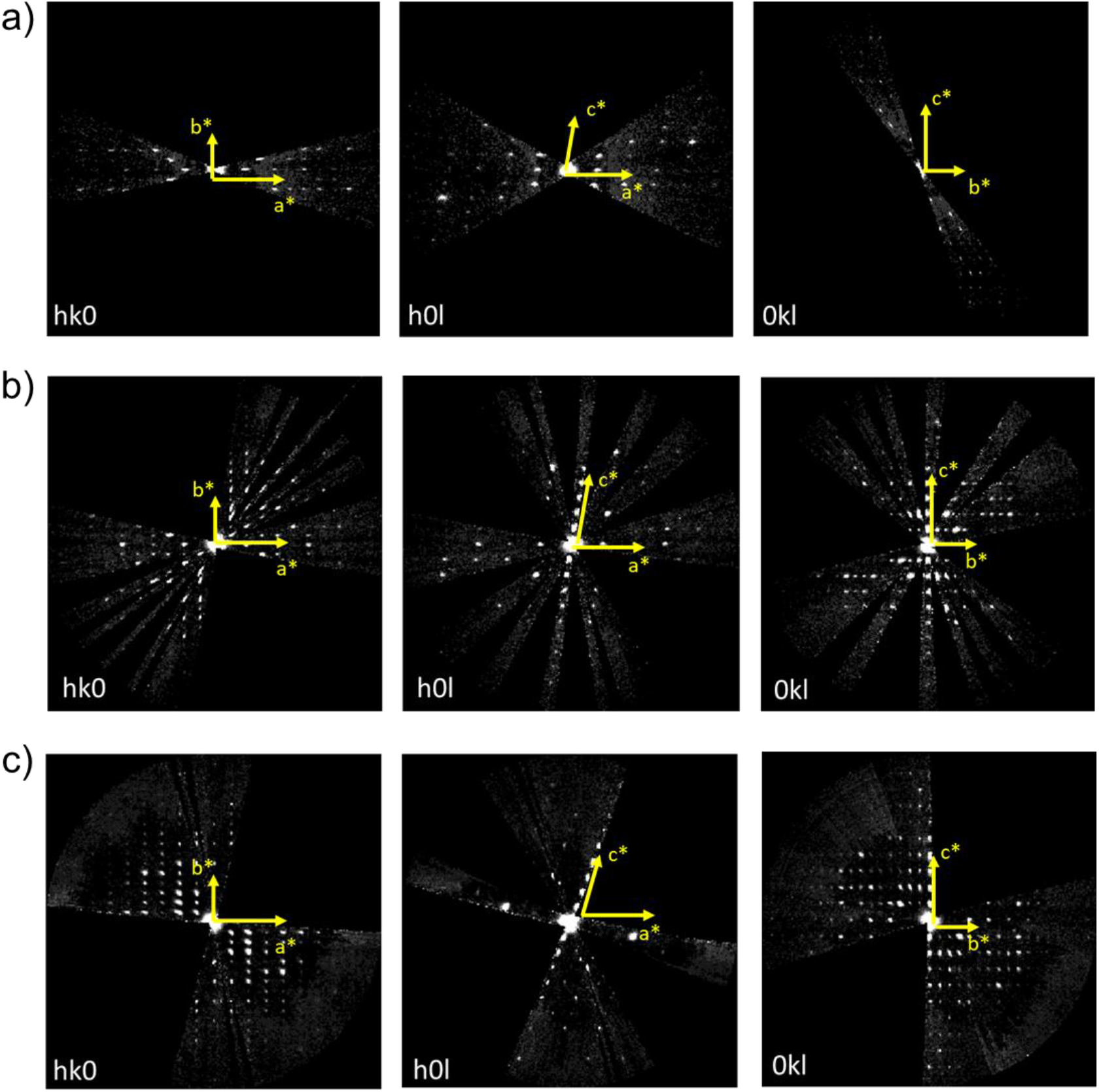
Images of the reciprocal sections projections hk0, h0l and 0kl respectively of phase I (a), phase II (b) and III (c).

**Figure S6.**
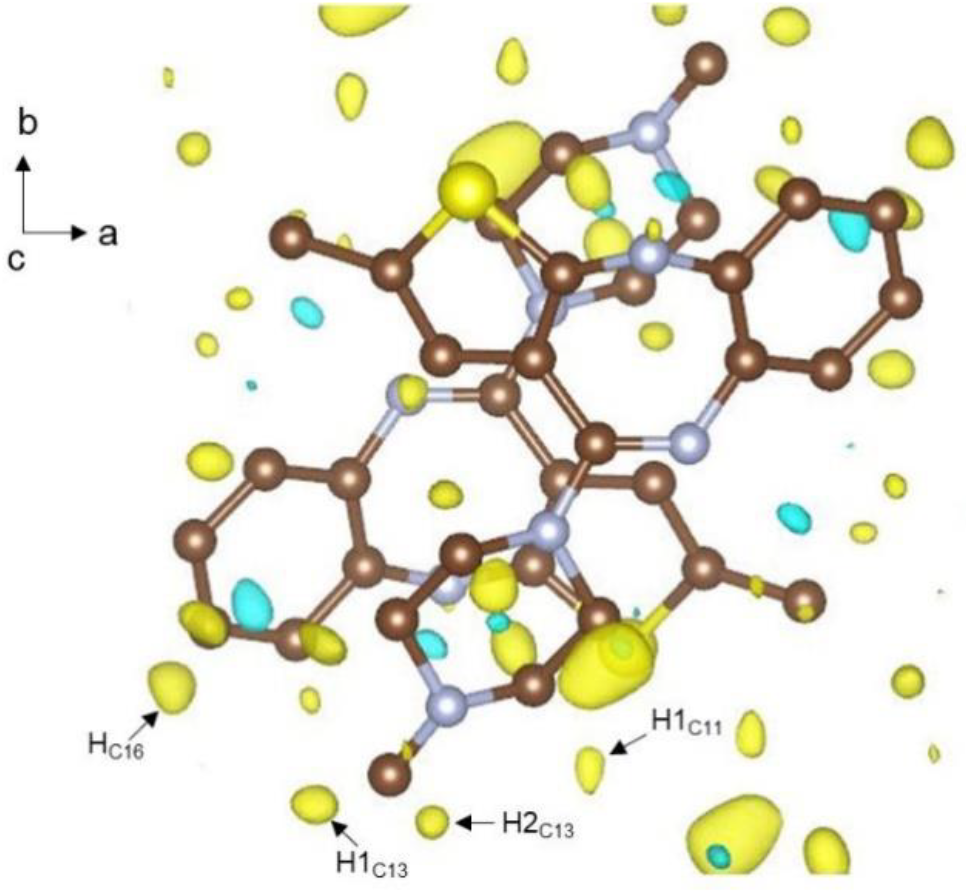
Difference Fourier map of OLZP form III before adding hydrogen atoms calculated at a 3σ isosurface level, just a few hydrogen atoms can be seen.

**Figure S7.**
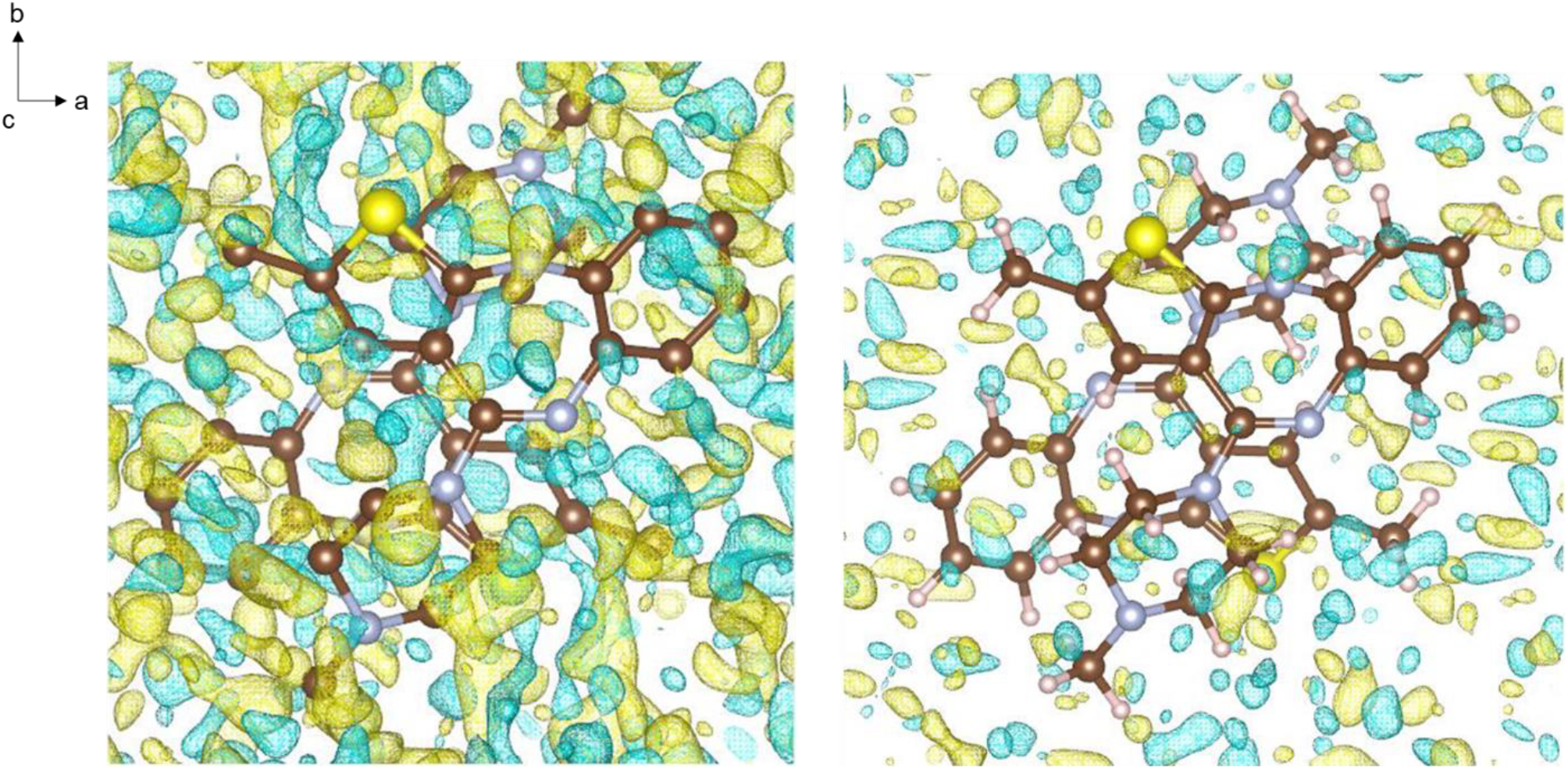
The two different Fourier maps of OLZP form III calculated at the same isosurface level before (left) and after (right) adding the expected hydrogen atoms viewed along the c-axis direction.

### Le Bail intensity extraction and Rietveld refinements results of the multiphase mixture

The unit cell parameters of forms I were not refined due to its little content. The refined cell dimensions after Le Bail extraction intensities are: for form II a = 9.933914(30) (Å), b = 16.54212(40) (Å), c = 10.01167(28) (Å), β = 98.06863(35)°; for the new form III they are a = 10.61363(19) (Å), b = 16.47721(28) (Å), c = 10.02741(26) (Å), β = 110.2544(21)°. The residuals factors of the fit are R_obs_ = 1.90%, wR_obs_ = 2.84%.

In the case of Rietveld refinement, the patterns were modelled with a mixture of forms III, II and I in infinitesimal quantity. The unit cell parameters of the latter were not refined. The refined cell dimensions and refinement parameters for form III and II are respectively a = 10.61457(34) Å, b= 16.47859(43) Å, c= 10.02385(30) Å, β = 110.2624(28)° and a = 9.93127(53) Å, b= 16.5275(66) Å, c= 10.01208(64) Å, β = 98.0086(57)°. The relative agreement factors are R_obs_ = 3.05% and wR_obs_ = 4.25% for form III, R_obs_ = 3.40% and wR_obs_ = 4.51% for form II, R_obs_ = 3.28% and wR_obs_ = 3.65% for form I.

**Figure S8.**
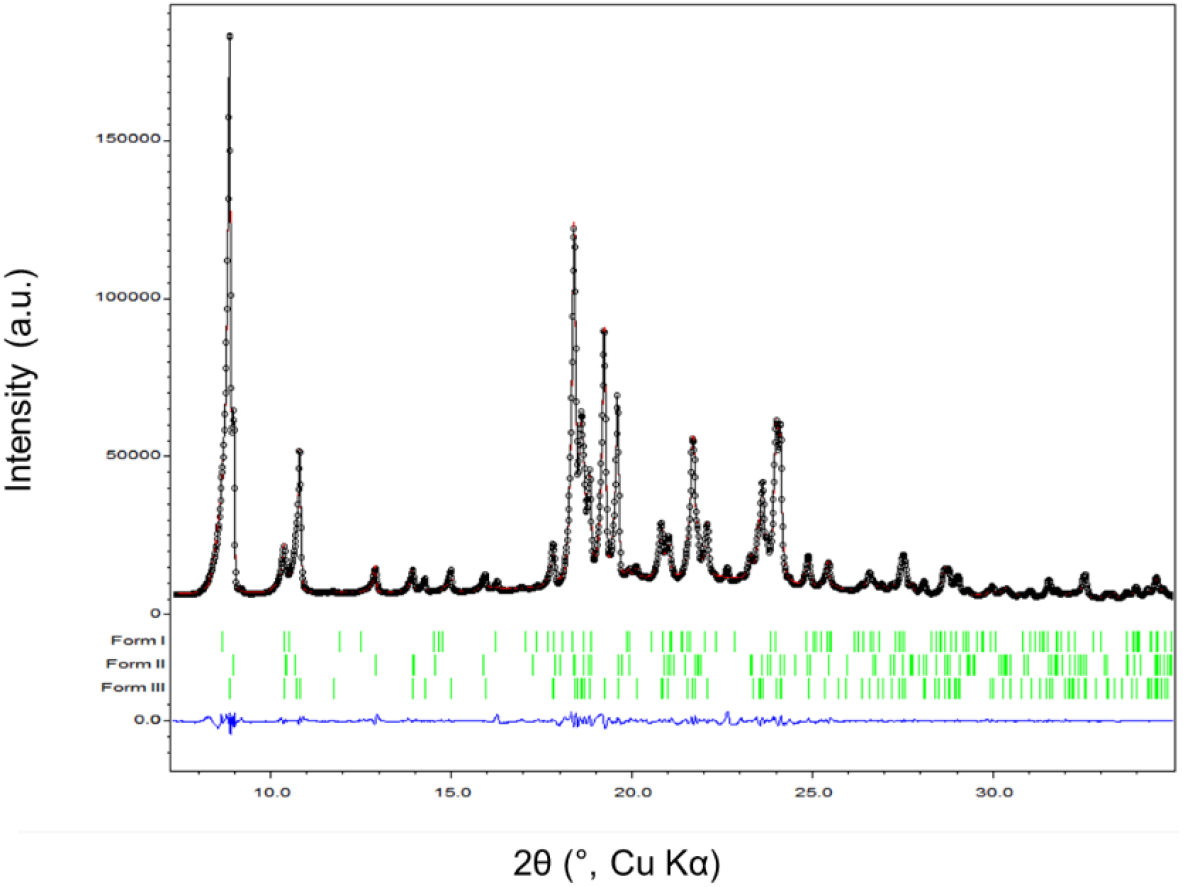
Le bail intensities extraction of the mixed forms I, II and III of OLZP obtained after recrystallisation.

#### 2D-finger prints of Hirshfeld surfaces of forms II and III

**Figure S9.**
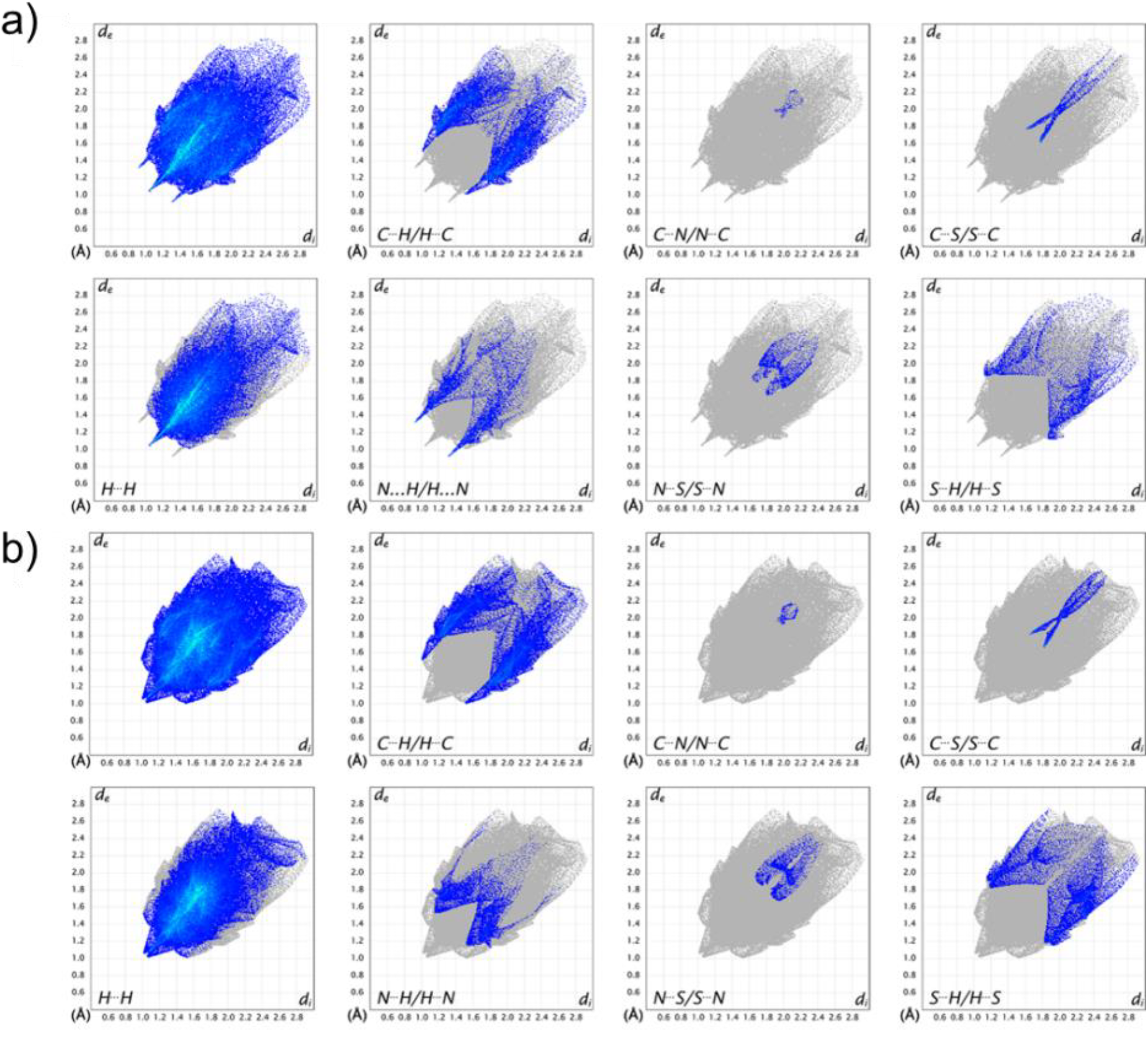
Comparison between fingerprint plots for the single molecule of OLZP in form II (A) and form III (B).

**Table S2.**
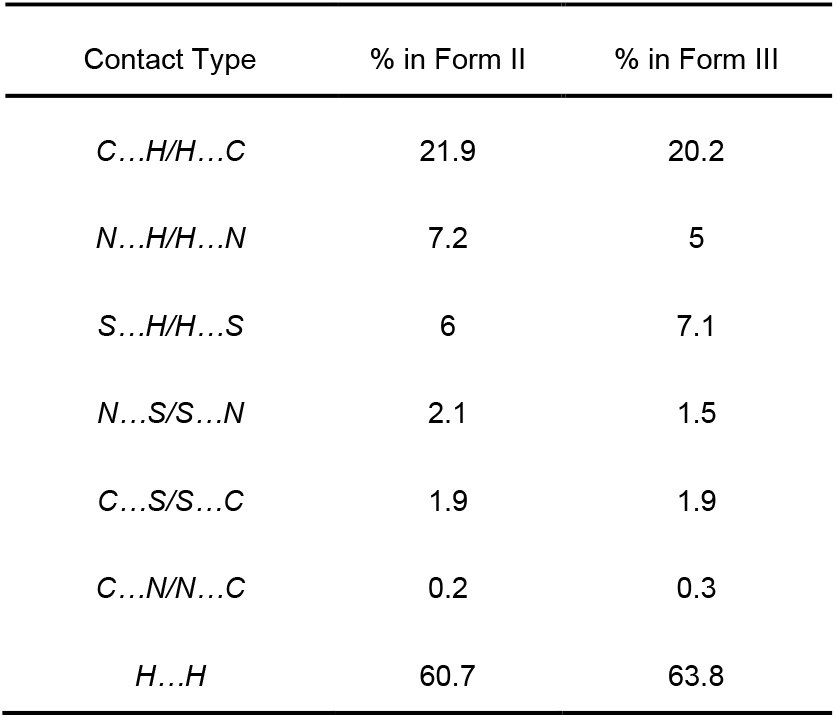
Percentage contributions to the Hirshfeld surface area of the different close intermolecular contacts of OLZP molecule in forms II and III.

### Comparison of A162 with form II

Figure S9a shows the packing of two layers (green and pink) of the structure A162 viewed along the relative b-axis direction. Hiding the pink layer and viewing the structure along [-1 2 0] direction what we see is the equivalent of the green layer encountered in forms II and III (fig. S9a). On the other hand, omitting the green layer and viewing the structure along the [1 2 0] direction (fig. S9c) the pink layer appears shifted of one half along the bc-plane, but the presence of the glide plane parallel to the (001) and normal to the (010) causes the exchange of the SC_0_ enantiomers every second layer.

**Figure S10.**
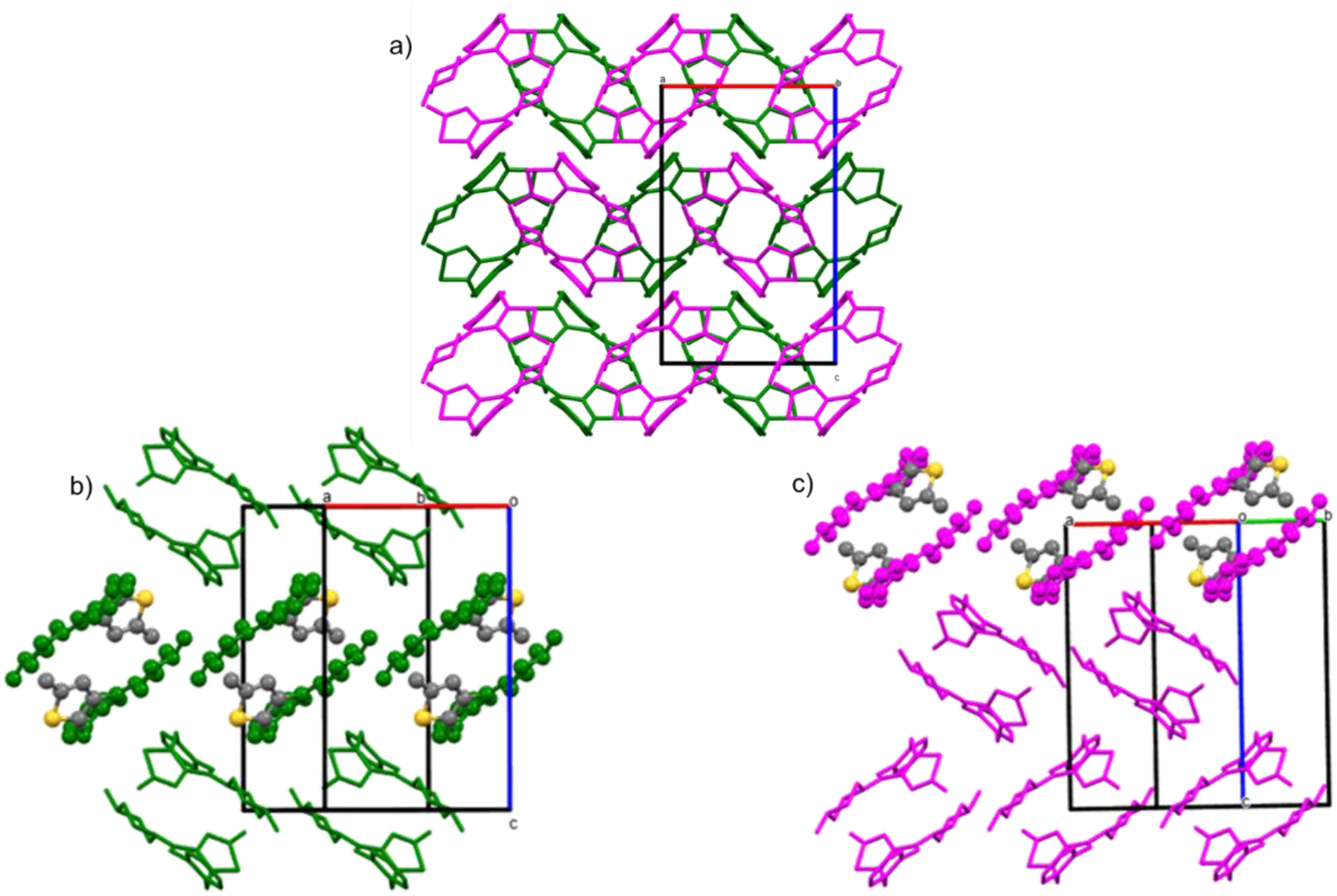
Packing of two layers in structure A162 viewed along the b-axis (a); green equivalent layer viewed along the [-1 2 0] direction (b) and the pink layer viewed along the [1 2 0] direction (c).

#### Comparison of UNOGIN_eq125 with phase III

**Figure S11.**
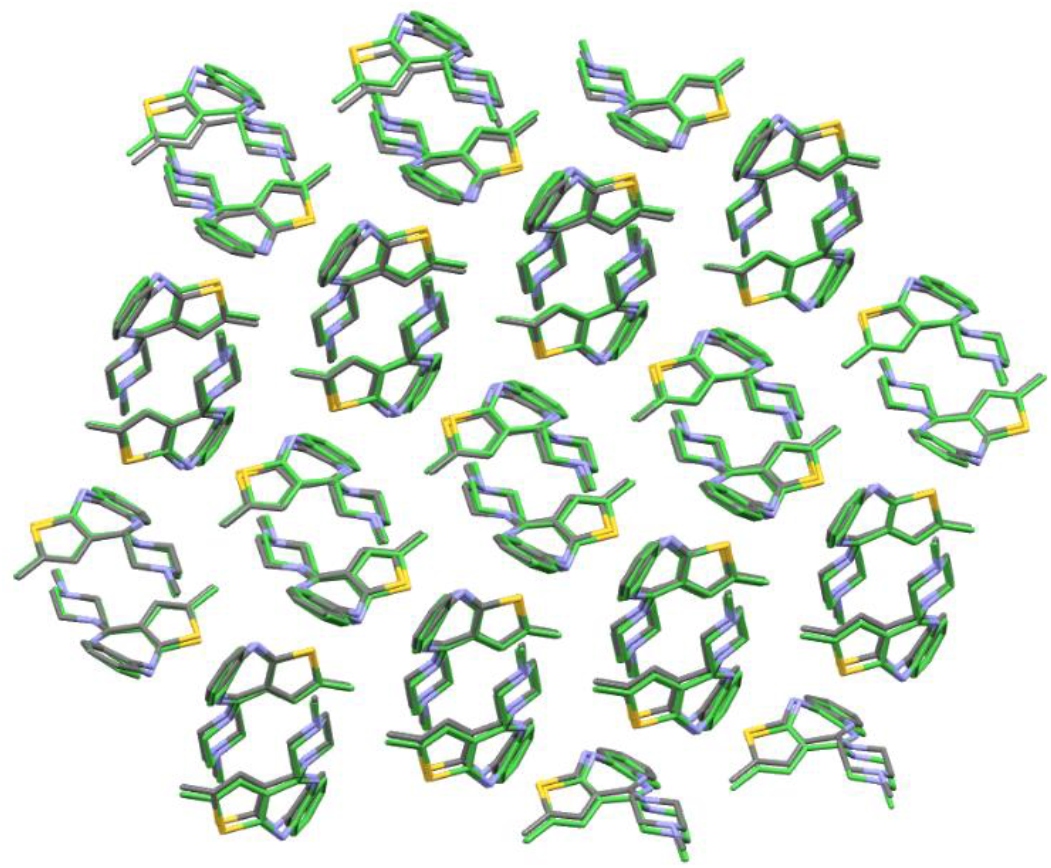
Overlay of the experimental form III (colored by element with C gray, N light blue, and S yellow) with UNOGIN_eq125 (green).

**Table S3.** Unit cell parameters of OLZP form III and the predicted structure UNOGIN_eq125.

| Polymorph | III | UNOGIN_eq125 |
| --- | --- | --- |
| a (Å) | 10.71(5) | 11.7 |
| b (Å) | 16.48(7) | 16.9 |
| c (Å) | 10.07(5) | 10.2 |
| $\alpha$ (°) | 90 | 90 |
| $\beta$ (°) | 110.4(2) | 58.4 |
| $\gamma$ (°) | 90 | 90 |
| V (Å<sup>3</sup>) | 1664 | 1717 |

**Figure S12.**
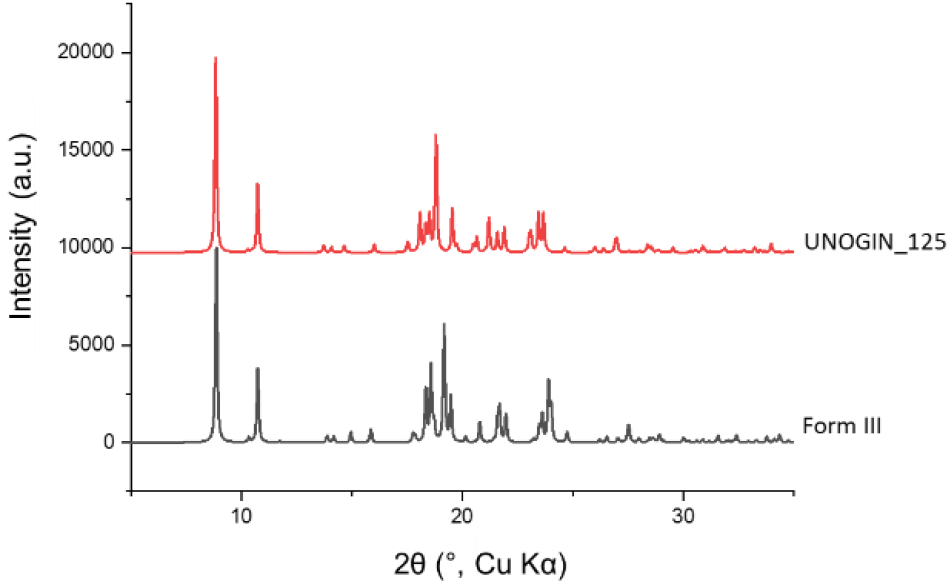
Comparison of the calculated PXRD patterns of phase III (black) with the predicted structure UNOGIN_eq125. The two graphs differ especially between the 2θ range of 15° and 25°.

### Accession Codes

CCDC 2309728 (deposition number of the crystal structure of olanzapine form III) contains the supplementary crystallographic data for this paper. These data are provided free of charge by the joint Cambridge Crystallographic Data Centre and Fachinformationszentrum

## Acknowledgements

The authors acknowledge Rajni M. Bhardwaj (University of Strathclyde) for supplying detailed information about olanzapine syntheses and Dr. Louise S. Price (UCL Chemistry) for computing crystal energy landscape and repository of predicted crystal structures and Ashwin Suresh for assistance in the use of Jana2020 for dynamical refinement.

